# The aggregate effect of implementation strength of family planning programs on modern contraceptive use at the health systems level in rural Malawi

**DOI:** 10.1101/2020.04.17.046524

**Authors:** Anooj Pattnaik, Diwakar Mohan, Amy Tsui, Sam Chipokosa, Hans Katengeza, Jameson Ndawala, Melissa A. Marx

## Abstract

**Background:** To explore the association between the strength of implementation of family planning (FP) programs on the use of modern contraceptives. Specifically, how strongly these programs are being implemented across a health facility’s catchment area in Malawi and the odds of a woman in that catchment area is using modern contraceptives. This information can be used to assess whether the combined impact of multiple large-scale FP programs is leading to change in the health outcomes they aim to improve.

**Methods and findings:** We used data from the 2017 Implementation Strength Assessment (ISA) that quantified how much of family planning programs at the health facility and community health worker levels were being implemented across every district of Malawi. We used a summary measure developed in a previous study that employs quantitative methods to combine data across FP domains and health system levels. We tested the association of this summary measure for implementation strength with household data from the 2015 Malawi Demographic Health Survey (DHS). We found that areas with stronger implementation of FP programs had higher odds of women using modern contraceptives compared with areas with weaker implementation. The association of ISA with use of modern contraception was different by education, marital status, and geography. After controlling for these factors, we found that the adjusted odds of using a modern contraceptive was three times higher in catchment areas with high implementation strength compared to those with lower strength.

**Conclusion:** Metrics that summarize how strongly FP programs are being implemented were used to show a statistically significantly positive relationship between increasing implementation strength and higher rates of modern contraceptive use. Decisionmakers at the various levels of health authority can use this type of summary measure to better understand the combined impact of their diverse FP programming and inform future programmatic and policy decisions. The findings also reinforce the idea that having a well-supported and supplied cadre of community health workers supplementing FP provision at the health facility can be an important health systems mechanism, especially in rural settings and to target youth populations.

## Introduction

An estimated 225 million women across the world have an unmet need for family planning (FP), and regions such as sub-Saharan Africa have much lower modern contraceptive prevalence rates (mCPR) and high fertility rates.^1,2,3^ It is dismaying that so many women are denied their fundamental human right to access the FP services and methods they desire.^4^ Increased use of these FP methods can have a tremendous economic benefit as well; for instance, helping to achieve the demographic dividend where more of the population are working rather than dependent, sparking much-desired economic growth in these low-income countries.^5,6^

The literature suggests that increasing the accessibility and readiness of the health system through programs that target the training and supervision of health workers, expanding method choice at service delivery points (SDPs), or demand generation activities for FP can have a positive impact on unmet need and mCPR among women of reproductive age.^7,8,9^ Other strategies include increasing the density (mean number of workers per population) of health workers providing FP, including task shifting FP delivery to community health workers (CHWs).^10,11^ Hence, many governments have chosen to design and implement large-scale family planning (FP) programs that include these activities in order to bolster the readiness and delivery of FP services across their health system.^12,13,14^

The sub-Saharan African country of Malawi is one of the poorest and youngest in the world, with nearly 70% of its 16.4 million population living below the international poverty line and 54% of this total population under 18.^15,16^ In the last decade, the country has experienced rapid population growth and high maternal mortality, which has increased their focus on implementing multiple national and subnational FP programs.^17,18,19^ This is evident in their recent national strategy policies and initiatives, such as their Youth-Friendly Health Services (YFHS) program.^20,21^

The Malawi FP delivery system includes government, faith-based (Christian Health Association of Malawi – “CHAM”), and NGO facilities, as well as two types of community health workers: paid Health Surveillance Agents (HSAs) who can provide male condoms, oral contraceptive pills (OCPs), and injectables (the most popular form of FP in Malawi) and voluntary Community-Based Distribution Agents (CBDAs), who can provide male condoms and OCPs directly to women.

Previous studies have shown mixed results in demonstrating a clear association between structural quality or readiness (an analogue to implementation strength) and contraceptive use.^22,23,24^ Yet, very few of these studies include outreach services or analyze the combined impact of the implementation of multiple FP programs at the national scale.^25,26,30^ We conducted an Implementation Strength Assessment (ISA) in 2017 to measure the strength of FP programs delivered across the community and facility levels in Malawi and developed a summary measure that reflects the combined implementation strength of FP programs across these levels.^27^ Using this summary measure, this paper reports on the association between the implementation strength of FP programs and modern contraceptive use among Malawian women.

## Materials and Methods

This study draws from two data sources from Malawi: the 2017 ISA of FP programs and the 2015 Malawi Demographic and Health Survey (DHS).

## Data source for independent variable

We reported previously on the methods the ISA conducted in Malawi in 2017.^27^ The ISA was a cross-sectional, mobile phone-based survey conducted from May to July 2017 that aimed to understand how strongly FP programs, especially those directed at youth, were being implemented at the health facility and CHW levels across all 28 districts of Malawi. Data were collected across five domains: training of health workers, supervision of health workers, contraceptive method and supply availability, FP demand generation, and accessibility of FP services.

## Creating an implementation strength score for each health facility’s catchment area

Creation of the composite IS score has been described in a previous study by Pattnaik et al.^28^ The IS score was created by (1) combining data across indicators (e.g. trained in the past year) and domains (e.g. training health workers); and (2) combining data across health facility (Facility In-Charge and Health Facility Worker) and community health workers (HSA and CBDA). The methods recommended in that study to combine indicator data, the principal components analysis (PCA), and to combine service provision level data, a mixed effects model (MEM), were used in this analysis. The scores resulting from PCA and the MEM were then adjusted by the density of health workers providing FP in that facility’s catchment area. The numerator for this density indicator is the numbers of facility and community health workers associated with each facility, data collected during the ISA. The denominator for the density indicator is each facility’s catchment area population from the 2008 Malawi census.^15^

In this study, the resulting summary measure represents the implementation strength of FP services at the catchment area level and was used as an independent variable in a regression with key FP outcomes from the 2015 Malawi DHS. We categorized the summary measure into quartiles for easier interpretation and comparison.

## Data Source for Outcome

Modern contraceptive use was chosen because it is the most proximate outcome indicator of the type of FP programs the ISA assessed and it is the main objective of recent MoH FP strategies and programs in Malawi.^20,21,29,30^ We used the DHS definition of this outcome variable, which is a female survey respondent reporting that she used any of male condoms, female condoms, oral contraceptive pills, injectables, implants, IUDs, male and female sterilization, or emergency contraception at the time of interview. The data source for modern contraceptive use was the 2015 Malawi DHS, which is a nationally representative survey that provides national, regional, urban/rural, and district estimates for household and respondent characteristics as well as key health statistics in Malawi.^18^ The following variables were also chosen from this DHS as possible confounders and effect modifiers of the relationship: age, education, marital status, region, religion, and economic status of individual women.^7^

## Linking facility catchment area implementation strength with individual-level FP outcomes collected in the DHS

We linked the population-level FP outcome data from the DHS with the implementation strength data at the facility catchment area level from the ISA using a catchment linking method proposed elsewhere but described briefly here.^31^ In DHS surveys that include HIV biomarkers, geo-coordinates of the rural enumeration areas (EA) clusters are displaced up to 5km from their original location in a random direction to protect participant confidentiality and 1% of these rural clusters are randomly selected and displaced up to 10 km away.^32^ To account for the uncertainty caused by this displacement, the catchment area method creates 5 km buffers around DHS cluster centroids and identifies the health facility catchment areas that fall within these buffers (Fig 1). We decided to exclude clusters designated as urban due to the multitude of health facilities within in an urban area. Rural clusters, however, had fewer facilities within them which helps to test the association between the implementation strength of a rural facility and women within the rural cluster who have far less choices in a health provider.

**Fig 1.**
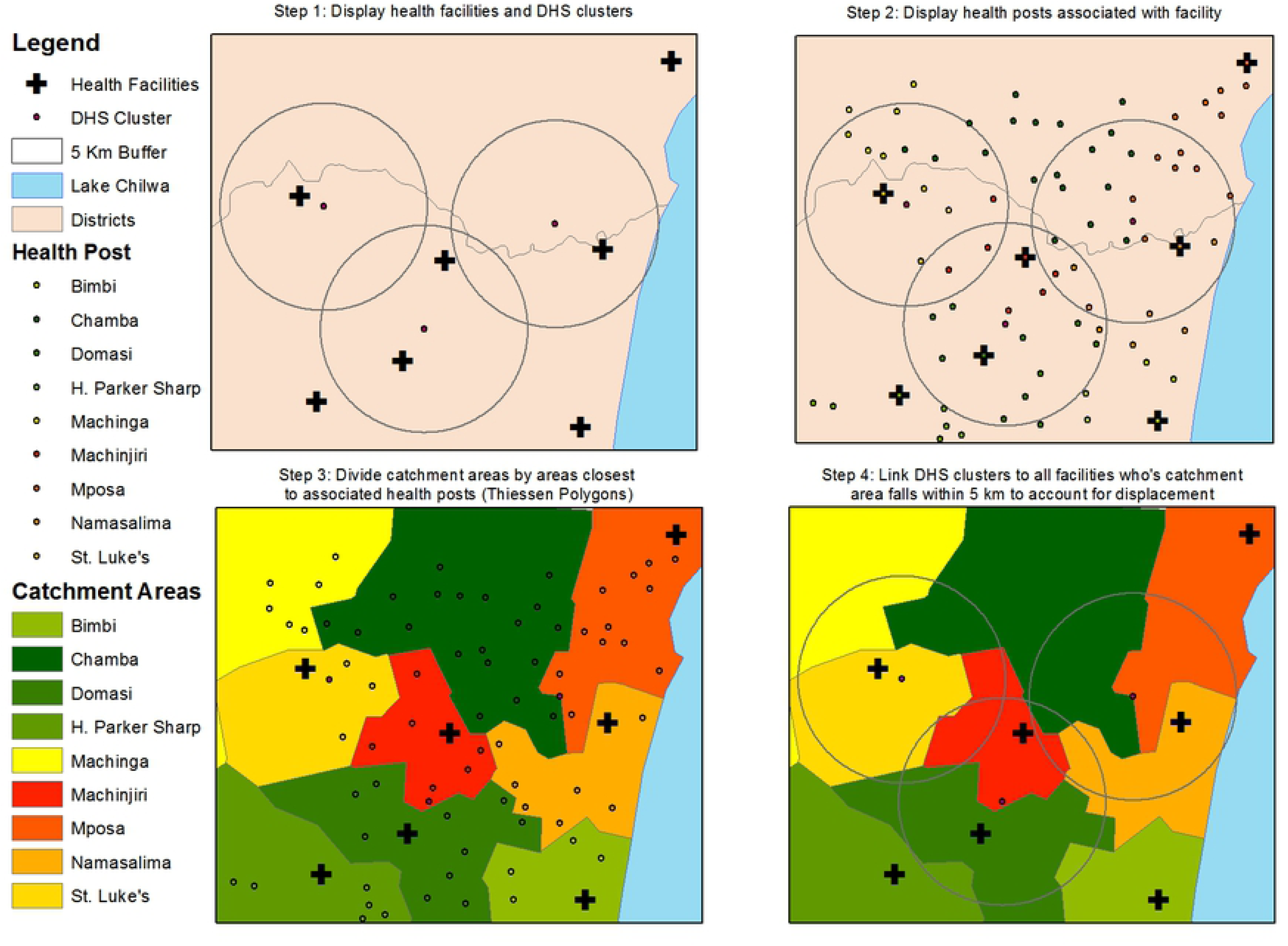
Steps to link health facility catchment area and population-level data via GIS^31^

Linked health facility catchment areas include the facility and community sites where CHWs provide FP, all of which contribute to the summary measure for implementation strength across a given catchment area.^28^ When these buffers captured the catchment areas of several facilities we averaged the IS scores across the DHS clusters for a single score. As a result, each DHS cluster (and data from the individual DHS respondents within them) was linked with a single IS score representing the facilities in that cluster. We excluded DHS clusters classified as urban because they have higher facility density and more transport options, thus can be an inconsistent predictor of service utilization in urban areas.^33^

## Analysis

We used a mixed effects model to test the association between the implementation strength of FP programs and modern contraceptive use among rural women at the catchment area level. This model allows us to analyze fixed and random effects, and to account for any clustering at the district level. The fixed effects in this model are the IS scores and the control variables listed earlier. The outcome is the odds of a woman currently using a modern contraceptive; thus, we estimated odds ratios (ORs) and 95% confidence intervals (CIs) in the regressions. Unadjusted and adjusted logistic regression models are compared. We used forward stepwise regression to build the model linking IS and modern contraceptive use, considering common confounders. We also tested models with interaction terms for the variables that were statistically significantly associated with our outcomes. All analysis was conducted using R 3.4.1 software.^34^

## Results

For the outcome data, we removed the urban clusters from the DHS which resulted in the dataset being comprised of 675 out of 850 total clusters (79%), with 19,261 out of 24,562 total women (78%) represented. For the exposure data, we interviewed 660 of 666 health facility In-Charges in Malawi using the 2017 Malawi ISA. We reached 1662 of 1815 (92%) HFWs, 4048 of 4131 (98%) HSAs selected, and 3187 of 3430 (93%) CBDAs for interview. Less than 10% of each HW cadre stated that they did not provide FP and less than one percent of those reached declined to participate. Chipokosa et al provides more detail of the data collected for the 2017 ISA.^27^ After restricting to rural facilities, 497 facilities were retained (Table 1).

**Table 1.** Background characteristics of rural health facilities

|  | n (%) |
| --- | --- |
| Total number of health facilities retained | 497 |
| Proportion of facilities that are health centers | 455 (91.5) |
| Managing authority of health facility |  |
| MoH | 362 (72.8) |
| CHAM* | 120 (24.1) |
| Other** | 15 (3.0) |
| Region (%) |  |
| North | 84 (16.9) |
| Central | 191 (38.4) |
| South | 222 (44.7) |
| Number of health workers per facility catchment area (median) |  |
| Health Facility Worker | 2 |
| Community Health Worker (HSA*** and CBDA****) | 11 |

\*CHAM: Christian Health Association of Malawi
\*\*NGO, police, or tea estate facilities
\*\*\*HSA: Health Surveillance Agent
\*\*\*\*CBDA: Community-Based Distribution Agent

Most (73%) rural facilities in this study are government health centers, with CHAM comprising another 24% of facilities. The Northern region has the lowest proportion of facilities (16.9%) compared to the other regions, which is commensurate with the region only comprising about 12% of the population.^15^ The facilities in this study have a median of two workers providing FP in the facility and 11 workers providing FP in the community. From this data, an IS summary measure was calculated for each facility’s catchment area (CA) or areas that comprise a DHS cluster (Fig 2).

**Fig 2.**
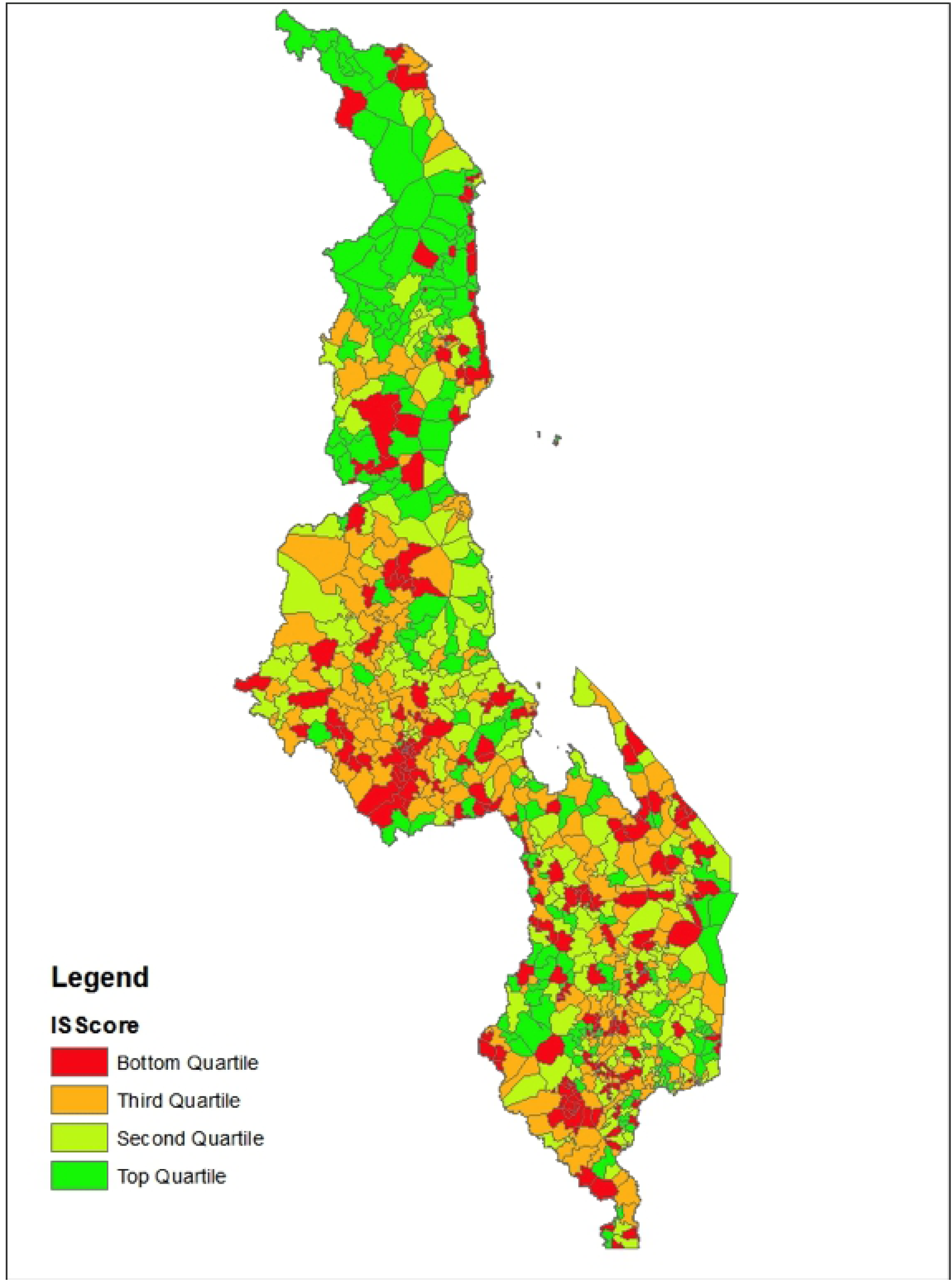
Heat map of IS scores within each DHS cluster across Malawi

More CAs in the northern region are in the top quartile of implementation strength, while the Central region appear to have more in the lowest quartile. Many CAs that are in the bottom two quartiles are in the Southern region.

Table 2 describes the characteristics of the 19,261 women who provided data within the matched rural clusters from the DHS. This table also lists the proportion of women using modern contraceptives stratified by socio-demographic characteristics.

**Table 2.**
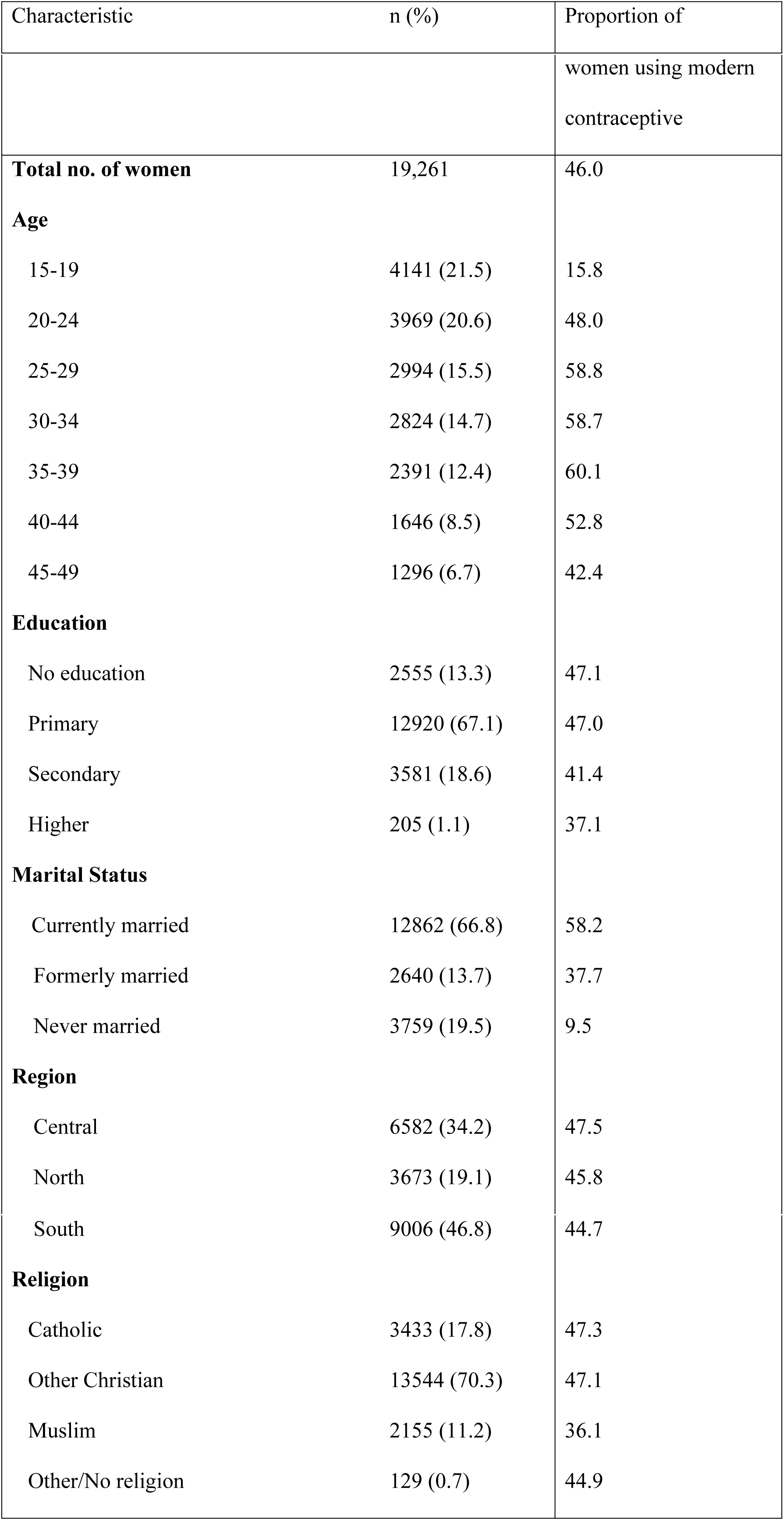
Background trait of women in rural DHS clusters and associated modern contraceptive use for each trait

| Characteristic | n (%) | Proportion of |
| --- | --- | --- |

|  |  | women using modern<br>contraceptive |
| --- | --- | --- |
| **Total no. of women** | 19,261 | 46.0 |
| **Age** |  |  |
| 15-19 | 4141 (21.5) | 15.8 |
| 20-24 | 3969 (20.6) | 48.0 |
| 25-29 | 2994 (15.5) | 58.8 |
| 30-34 | 2824 (14.7) | 58.7 |
| 35-39 | 2391 (12.4) | 60.1 |
| 40-44 | 1646 (8.5) | 52.8 |
| 45-49 | 1296 (6.7) | 42.4 |
| **Education** |  |  |
| No education | 2555 (13.3) | 47.1 |
| Primary | 12920 (67.1) | 47.0 |
| Secondary | 3581 (18.6) | 41.4 |
| Higher | 205 (1.1) | 37.1 |
| **Marital Status** |  |  |
| Currently married | 12862 (66.8) | 58.2 |
| Formerly married | 2640 (13.7) | 37.7 |
| Never married | 3759 (19.5) | 9.5 |
| **Region** |  |  |
| Central | 6582 (34.2) | 47.5 |
| North | 3673 (19.1) | 45.8 |
| South | 9006 (46.8) | 44.7 |
| **Religion** |  |  |
| Catholic | 3433 (17.8) | 47.3 |
| Other Christian | 13544 (70.3) | 47.1 |
| Muslim | 2155 (11.2) | 36.1 |
| Other/No religion | 129 (0.7) | 44.9 |

Nearly 67% of women reported being currently married and a much higher proportion of them reported using modern contraceptives compared to women who are not married. About 67% of women have only a primary education, which also has the highest proportion of modern contraceptive use in this category. Over 40% of rural women in Malawi are under the age of 25 and modern contraceptive use is substantially lower among those who are 15-19 years-old compared to women who are 20-24. Wealth distributions were not included in this table because there weren’t large differences between the quintiles.

Women had higher adjusted odds of using a modern contraceptive with higher catchment IS scores, in a dose-response relationship, however, with large confidence intervals (Table 3).

**Table 3.**
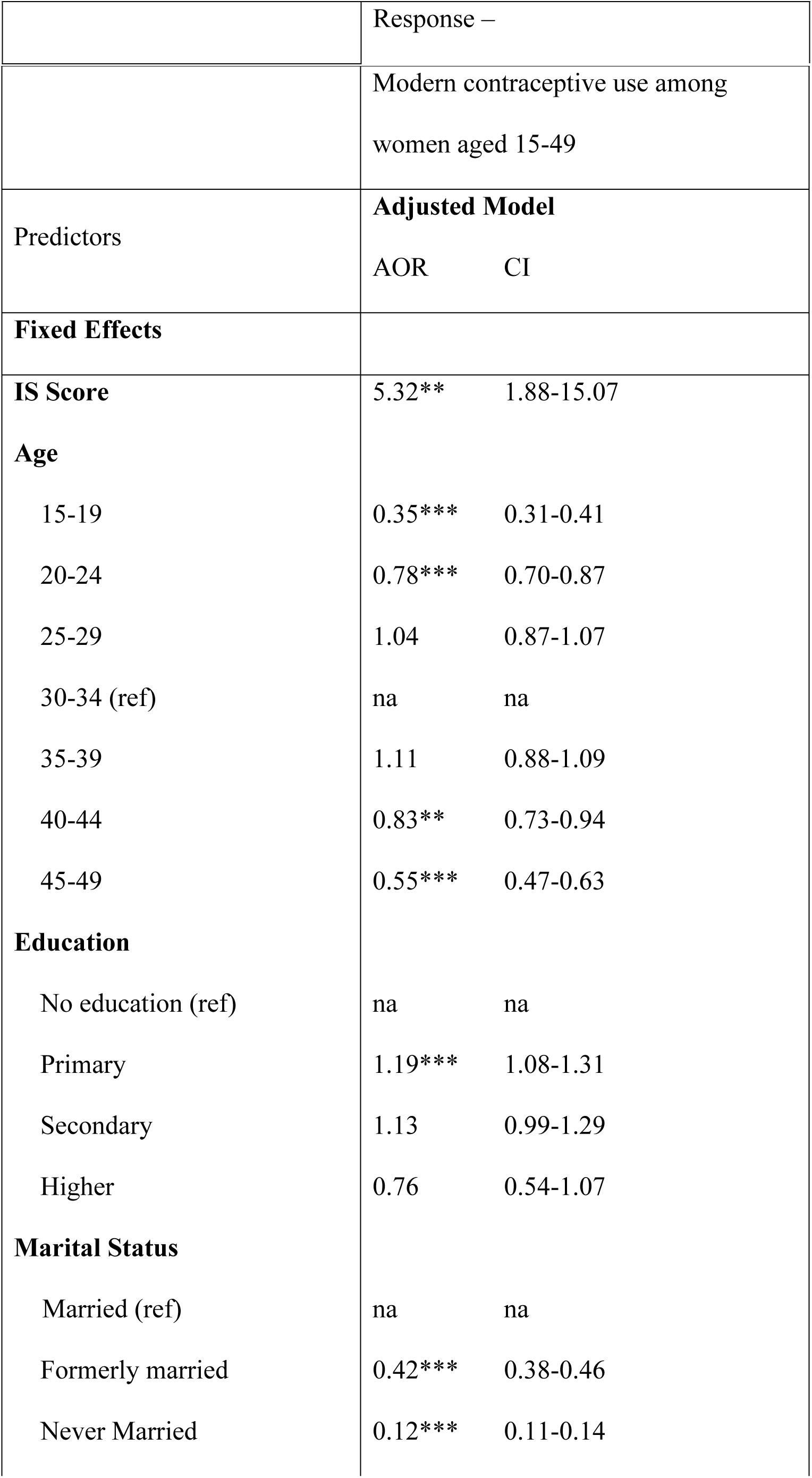
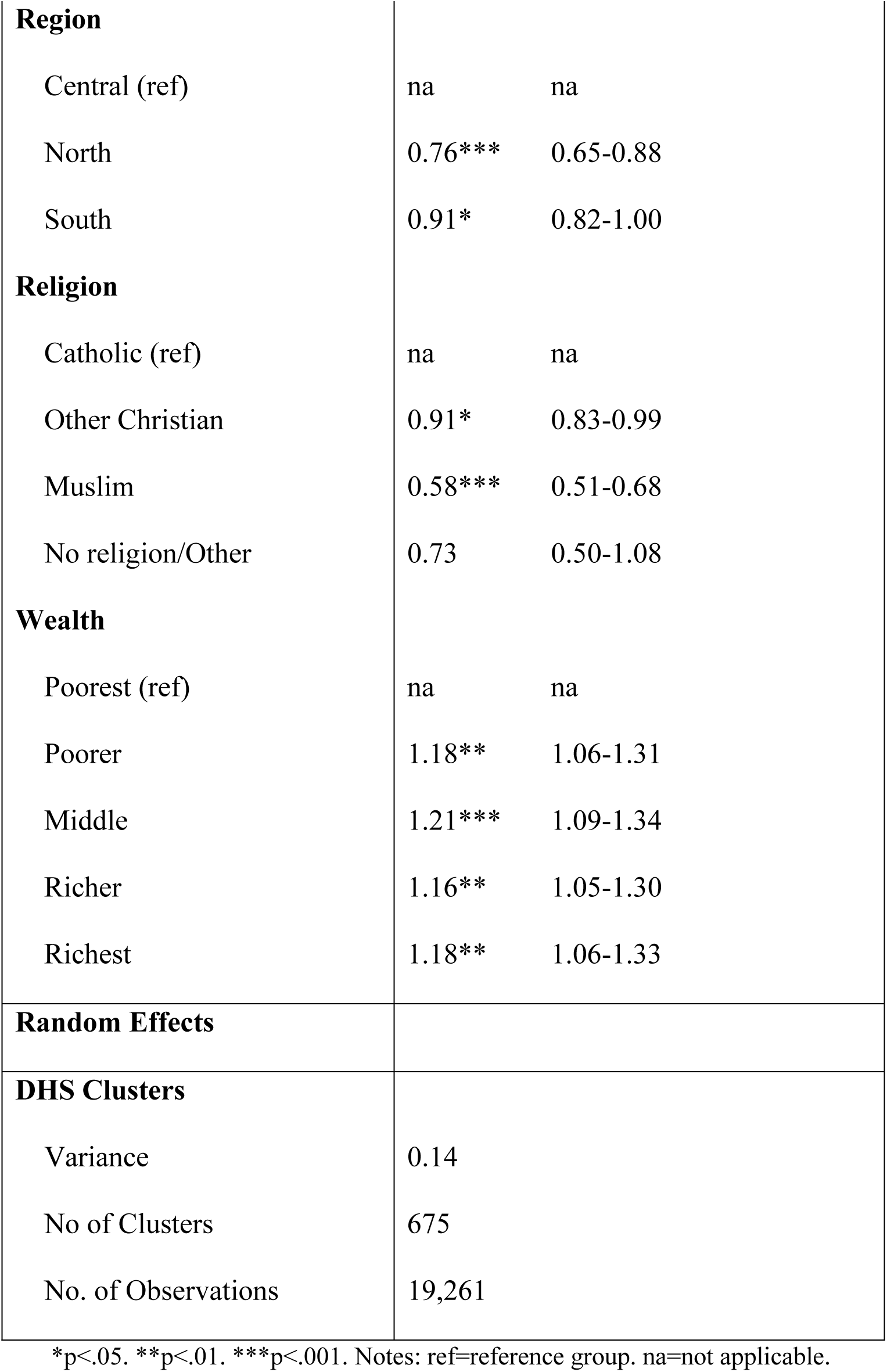
Association between rural women’s use of modern contraceptives and implementation strength score, adjusted model with random effects

|  |  |  |
| --- | --- | --- |
|  | Modern contraceptive use among women aged 15-49 |  |
| Predictors | **Adjusted Model** |  |
|  | AOR | CI |
| **Fixed Effects** |  |  |
| **IS Score** | 5.32** | 1.88-15.07 |
| **Age** |  |  |
| 15-19 | 0.35*** | 0.31-0.41 |
| 20-24 | 0.78*** | 0.70-0.87 |
| 25-29 | 1.04 | 0.87-1.07 |
| 30-34 (ref) | na | na |
| 35-39 | 1.11 | 0.88-1.09 |
| 40-44 | 0.83** | 0.73-0.94 |
| 45-49 | 0.55*** | 0.47-0.63 |
| **Education** |  |  |
| No education (ref) | na | na |
| Primary | 1.19*** | 1.08-1.31 |
| Secondary | 1.13 | 0.99-1.29 |
| Higher | 0.76 | 0.54-1.07 |
| **Marital Status** |  |  |
| Married (ref) | na | na |
| Formerly married | 0.42*** | 0.38-0.46 |
| Never Married | 0.12*** | 0.11-0.14 |
| **Region** |  |  |
| Central (ref) | na | na |
| North | 0.76*** | 0.65-0.88 |
| South | 0.91* | 0.82-1.00 |
| **Religion** |  |  |
| Catholic (ref) | na | na |
| Other Christian | 0.91* | 0.83-0.99 |
| Muslim | 0.58*** | 0.51-0.68 |
| No religion/Other | 0.73 | 0.50-1.08 |
| **Wealth** |  |  |
| Poorest (ref) | na | na |
| Poorer | 1.18** | 1.06-1.31 |
| Middle | 1.21*** | 1.09-1.34 |
| Richer | 1.16** | 1.05-1.30 |
| Richest | 1.18** | 1.06-1.33 |
| **Random Effects** |  |  |
| **DHS Clusters** |  |  |
| Variance | 0.14 |  |
| No of Clusters | 675 |  |
| No. of Observations | 19,261 |  |

\*p<.05. \*\*p<.01. \*\*\*p<.001. Notes: ref=reference group. na=not applicable.

Women who had primary (1.19***) or secondary education (1.13) had higher odds of using a modern contraceptive than women who had no education, although women who had the highest education level had lower odds (0.76, p<0.001) than the reference group. Further analysis found that education does not modify the effect between implementation strength and modern contraceptive use among women.

In comparison to the reference group of Catholic women, Muslim (0.58**, p<.05) and non-Catholic Christians (0.91*) had lower adjusted odds of using modern contraceptives. The relationship between implementation strength and modern contraceptive use among rural women did not significantly differ among the different religious groups measured.

For age, we used the 30-34 age group as the reference because this had the largest, most stable population with which to compare the other groups. Compared to 30-34 year-olds, and controlling for other factors, the adjusted odds of using modern contraceptives was statistically significantly lower for women of who are 15-19 (Adj OR=0.35***, CI=0.31-0.41), 20-24 (Adj OR=0.78***, CI=0.70-0.87), 40-44 (Adj OR=0.83**, CI=0.73-0.94), and 45-49 (Adj OR=0.55***, CI=0.47-0.63). The adjusted odds of women who were formerly married (0.42***, CI=0.38-0.46) or never married (0.12***, CI=0.11-0.14) using a modern contraceptive were statistically significantly lower than those who were married. The relationship between the implementation strength summary measures at the CA level and modern contraceptive use among rural women seems to differ between all seven age groups in the DHS (Fig 3).

**Fig 3.**
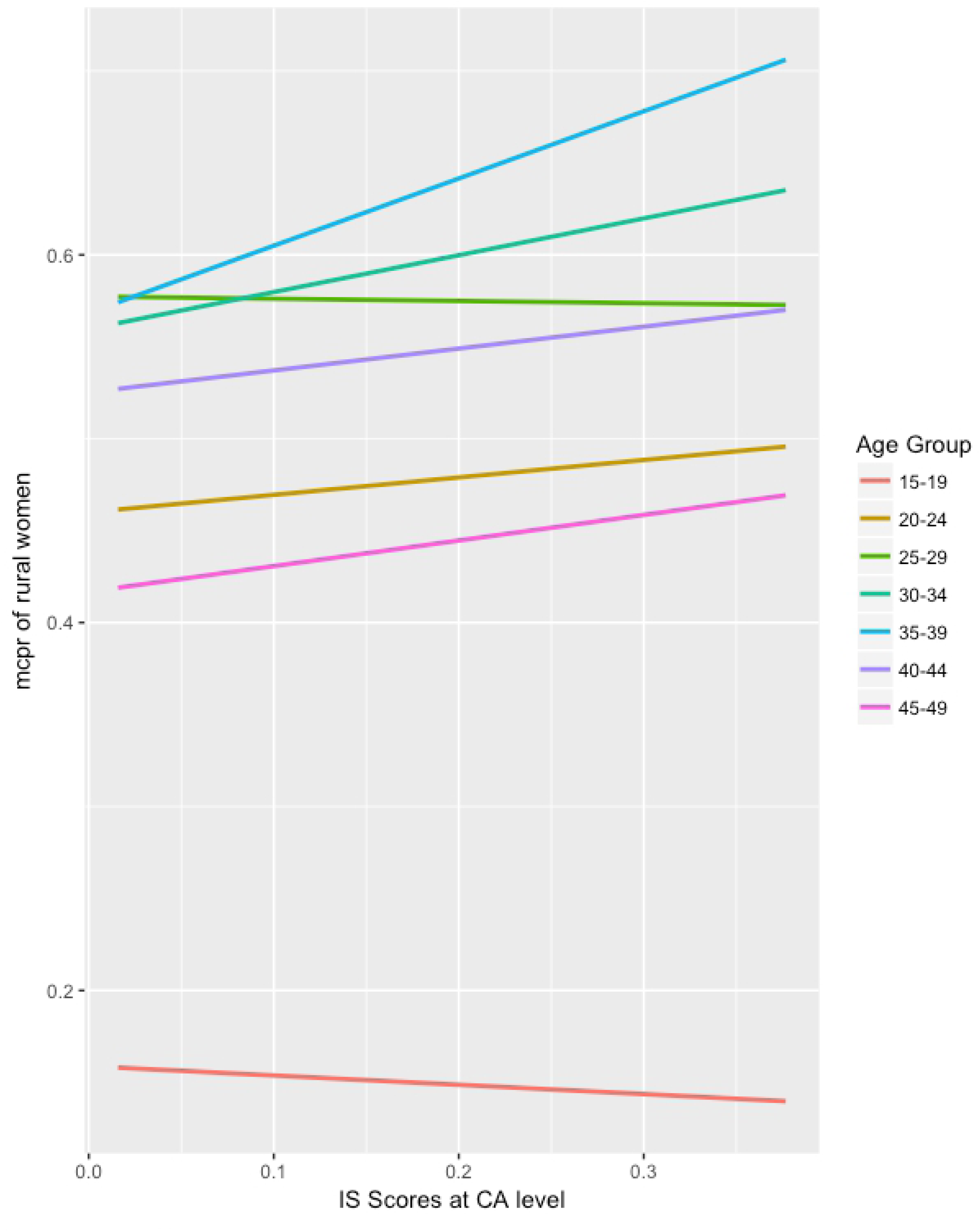
Relationship between implementation strength summary measure and modern contraceptive use among rural women, by age group

In the 15-19 and 25-29 age groups, there seems to be a slightly negative relationship between contraceptive use and implementation strength, whereas among all other women of reproductive age, stronger implementation seems to have a positive relationship with modern contraceptive use. However, none of the age groups were significant when tested for effect modification.

## Discussion

We found that areas with stronger implementation of FP programs had higher adjusted odds of women using modern contraceptives compared with areas with weaker implementation. The confidence intervals for the IS summary measures were larger than the other variables because these measures come from the smaller, ISA dataset. The association of implementation strength with use of modern contraceptives may be different by education, marriage, and region, which is consistent with recent literature that finds that increasing education among especially rural women has a positive effect on demand and mCPR.^29,35^ This finding adds to the debate about whether FP programs and services or development investments (such as education) are driving changes in contraceptive use and fertility.^25,36,37,38^ Similarly, cultural norms about marriage likely play a major role in the contraceptive decisions among these women,^25,29^ After controlling for these factors, we found that the adjusted odds of a woman using a modern contraceptive was three times as high in areas with strong implementation compared to those with weaker implementation.

There have been mixed results among studies aiming to demonstrate a link between structural quality (an analogue to IS) and contraceptive use.^23,24, 25,29,30,39,40^ Most of these studies are restricted to health facility data (often from Service Provision Assessments) and do not take outreach services into account like this study did. This is crucial to understand better, as task shifting FP provision to CHWs is a cost-efficient strategy that health authorities are increasingly employing. Additionally, this study fills a gap by analyzing how the combined strength of several FP programs being implemented can have an impact on modern contraceptive use. Most other studies we reviewed evaluate either single, specific programs or the readiness of health facilities (either without including outreach or analyzing it separately).^29,30,40^

While age didn’t seem to be a driver of increased modern contraceptive use, there were significant differences in the relationship between rural women of different reproductive age and their likelihood to use modern contraceptives. Stronger implementation of FP programs was a more important factor among 30-34 year-olds on whether they use modern contraceptives than any other age groups. One of the reasons that account for this is women at this age in Malawi likely have had at least one child and are more comfortable seeking modern contraceptives from the formal health system, without fear of stigma like the lower age groups.

While stronger implementation seemed to have a positive relationship with modern contraceptive use in most age groups, this relationship did not seem to be the same in the 15-19 age group.

These youth are the explicit target of government programs such as the Youth-Friendly Health Services (YFHS) in Malawi. In fact, there seemed to be an inverse relationship between implementation strength and odds of using a modern contraceptive in this group, which could suggest that these programs may have little impact on the youth or conversely, could point to the fact that youth often underreport their contraceptive use. This finding is consistent with other studies that point to demographic traits and cultural norms (such as a young marriage age) being major drivers of improvements in mCPR.^25,33,41^ Additionally, Malawian youth may use pharmacies, shops, and informal outlets rather than the formal health system to obtain their contraceptive methods, especially considering popular methods among youth are short acting ones like male condoms. The other group that the YFHS programs targets, 20 to 24-year-olds, are more likely to be married and thus, more likely to access modern contraceptives from the formal health system that our ISA measures.

Another unique aspect of this study is its use of the linking method to make maximum use of the data available in the 2017 Malawi ISA. Previous literature, such as Skiles et al and Peters et al, explored many different methods that can link service environment and population estimate data.^31,33^ We used the recommended linking method from the Peters et al study, which suggests this method was a better fit especially given the datasets used in this study.^31^ This method allowed us to analyze linkages at a more granular level than the facility or district: the catchment area level and among individual women in DHS clusters. These methods could be used by national or subnational authorities to better understand the aggregate effect of multiple FP programs being implemented in their jurisdictions on specific outcomes they seek to influence. It also allows them to better understand the nature of this relationship, how it affects different segments of the population, and can inform data-driven adjustments at the policy and programmatic levels. Future studies could explore using this method to test the association between implementation strength and population-level outcomes in other contexts and in other topic areas outside of FP.

## Limitations

We were limited by the data sources available for this study. We cannot claim causality because we used the 2015 Malawi DHS and the 2017 Malawi ISA; the outcome preceding the exposure. We assumed that FP outcomes did not change dramatically from the time the DHS was collected to when the IS data were collected. Thus, if IS became stronger since the DHS data were collected, then the association may be underestimated because the effects hadn’t been measured yet. Still, IS measured in 2017 could reflect the implementation of programs from a number of years before, thus being closer to the time of DHS data collection. Moreover, the cross-sectional nature of both the ISA and DHS datasets and the lack of counterfactual never allowed for statements about causation in the first place. We suggest that this analysis should be done again when the 2020 DHS in Malawi is conducted and released.

The 2008 Malawi census was used to generate HW density. Therefore, even in a country with a recognized health worker shortage for several years, density is likely overestimated.^42^ We recommend recalculating HW density when the 2018 Malawi census data are released. We also could not include NGO facilities in this analysis because we did not have their CA populations to calculate HW density. Although the remaining MoH and CHAM facilities comprise nearly 90% of all facilities in Malawi, we acknowledge missing these non-public facilities. Jayachandran et al (2016) found that NGOs have higher quality of care than government ones in Malawi and this could have a differential impact on youth populations.^43^ Future studies could target the association between the strength of implementation at NGO facilities and key FP outcomes.

Future research could also explore the connection between the structural indicators of the ISA and more process-oriented indicators (such as quality of care) that likely lie between IS and modern contraceptive use. Measuring provider-client interactions could shed light on how much implementation strength of FP programs in Malawi is affecting the experiences of clients on the ground. Along this chain, it would also be interesting to look at the link between implementation strength, modern contraceptive use, and fertility rates. While mCPR has been increasing consistently in Malawi over the last few decades, the total fertility rate (TFR) has stagnated recently. There is likely a number of reasons for this, including the high use of short-term methods, early age of marriage and first birth.^25,44^ Future research is needed to explore how the implementation strength of FP programs has an effect down the causal pathway, especially with more timely datasets.

## Conclusions

We developed a metric to quantifies the implementation strength of FP programs in Malawi and using this metric, we found that stronger implementation was associated with higher odds of modern contraceptive use. The data also suggests that this relationship may not be as strong with younger women and alternative approaches to target these populations likely should be explored. Especially in lower income countries where donor-led programs are rampant and often uncoordinated, government authorities at the national and subnational levels can use this metric to better understand the combined impact of these programs on the populations they target. This knowledge can they be used as a basis for future programmatic and policy adjustments, as well as in negotiations with external funders and implementers. Moreover, the findings suggest that in order to address population issues in low-resource settings and potentially reach the demographic dividend, it is not only important to strongly implement a variety of FP programs at the facility level, but to also supplement this with a well-prepared cadre of community health workers who can provide further outreach to rural women.

## Acknowledgments

The authors would also like to thank the following organizations and individuals for their valuable support of this work: the RHD Unit of the Malawi Ministry of Health, including Fannie Kachale; the Malawi National Statistics Office, including Mercy Kanyuka, Sautso Wachepa, and Lewis Gombwa; the Institute for International Programs (IIP) including Neff Walker, Agbessi Amouzou, Tim Robertson, and Emily Wilson; and Scott Zeger, Scott Radloff, Olakunle Alonge at the Johns Hopkins Bloomberg School of Public Health.

